# DISRUPTION OF TRANSTHALAMIC CIRCUITRY FROM PRIMARY VISUAL CORTEX IMPAIRS VISUAL DISCRIMINATION IN MICE

**DOI:** 10.1101/2025.02.07.637190

**Authors:** C. McKinnon, C. Mo, S. M. Sherman

## Abstract

Layer 5 (L5) of the cortex provides strong driving input to higher-order thalamic nuclei, such as the pulvinar in the visual system, forming the basis of cortico-thalamo-cortical (transthalamic) circuits. These circuits provide a communication route between cortical areas in parallel to direct corticocortical connections, but their specific role in perception and behavior remains unclear. Using targeted optogenetic inhibition in mice performing a visual discrimination task, we selectively suppressed the corticothalamic input from L5 cells in primary visual cortex (V1) at their terminals in pulvinar. This suppresses transthalamic circuits from V1; furthermore, any effect on direct corticocortical projections and local V1 circuitry would thus result from transthalamic inputs (e.g., V1 to pulvinar back to V1 (Miller-Hansen and Sherman, 2022). Such suppression of transthalamic processing during visual stimulus presentation of drifting gratings significantly impaired discrimination performance across different orientations. The impact on behavior was specific to the portion of visual space that retinotopically coincided with the V1 L5 corticothalamic inhibition. These results highlight the importance of incorporating L5-initiated transthalamic circuits into cortical processing frameworks, particularly those addressing how the hierarchical propagation of sensory signals supports perceptual decision-making.

**SIGNIFICANCE STATEMENT:** Appreciation of pathways for transthalamic communication between cortical areas, organized in parallel with direct connections, has transformed our thinking about cortical functioning writ large. Studies of transthalamic pathways initially concentrated on their anatomy and physiology, but there has been a shift towards understanding their importance to cognitive behavior. Here, we have used an optogenetic approach in mice to selectively inhibit the transthalamic pathway from primary visual cortex to other cortical areas and back to itself. We find that such inhibition degrades the animals’ ability to discriminate, showing for the first time that specific inhibition of visual transthalamic circuitry reduces visual discrimination. This causal data adds to the growing evidence for the importance of transthalamic signaling in perceptual processing.

## INTRODUCTION

Sensory processing relies heavily on bidirectional communication between thalamus and cortex (Sherman and Guillery, 2013; Usrey and Sherman, 2021). This includes projections from cortical layer 5 (L5) that robustly activate higher-order (HO) thalamic neurons (Sherman and Guillery, 2013; Usrey and Sherman, 2019; Sherman and Usrey, 2024) and initiate cortico-thalamo-cortical (transthalamic) circuits (Sherman and Guillery, 1998; Theyel et al., 2010; Sherman and Guillery, 2013; Sherman, 2016; Mo and Sherman, 2019; Usrey and Sherman, 2021; Miller-Hansen and Sherman, 2022; Mo et al., 2024; Sherman and Usrey, 2024). These transthalamic pathways run parallel to direct corticocortical projections. The appreciation of transthalamic processing has produced a transformative revision to conventional theories of cortical processing (Aru et al., 2019; Wolff et al., 2021; Suzuki et al., 2023).

Visual discrimination behavior, for example, relies on communication between primary (V1) and higher visual cortical areas (Jin and Glickfeld, 2020; Javadzadeh and Hofer, 2022), raising questions regarding the role of transthalamic circuitry in such behavior (Blot et al., 2021). Until recently, such causal influence of one cortical area on another has been attributed solely to direct corticocortical projections, but suppression of V1 also disrupts its transthalamic contribution, which was unappreciated. Thus, more selective functional circuit dissection is necessary to identify transthalamic contribution to behavior.

Indeed, V1 L5 provides the primary driving input to the pulvinar [formerly called the lateral posterior nucleus in rodents (Zhou et al., 2017)], a HO thalamic nucleus. Silencing V1 greatly diminishes visual responses in much of pulvinar (Harting et al., 1972; Bender, 1983; Casanova, 1993; Baldwin et al., 2017; Zhou et al., 2017; Blot et al., 2021; Kirchgessner et al., 2021), and recent findings indicate that L5 input specifically underlies this effect (Kirchgessner et al., 2021; Miller-Hansen and Sherman, 2022). Although L5 corticothalamic inputs initiate transthalamic circuitry, few *in vivo* studies have examined the function of these L5 inputs to pulvinar. One such study showed that inhibiting V1 layer 6 (L6) input to the lateral geniculate nucleus or pulvinar had no appreciable effect on visual responses of target thalamic cells. This presumably results because L6 provides relatively weak (modulator) input, whereas V1 L5 provides much stronger (driver) input to thalamus and thus is needed for normal responsiveness of pulvinar cells (Sherman and Guillery, 2013; Kirchgessner et al., 2021). However, most research on functional relationships between pulvinar and cortex has focused on thalamocortical rather than corticothalamic circuitry (Purushothaman et al., 2012; Casanova and Chalupa, 2023), showing that pulvinar inputs to cortex play a key role in integrating sensory input within the broader behavioral context (Roth et al., 2016; Blot et al., 2021), including coordinating the transmission of information between cortical areas in accordance with the salience of sensory stimuli (Saalmann et al., 2012; Zhou et al., 2016). Yet inhibiting the somatosensory L5 corticothalamic projection during a whisker-based perceptual task significantly disrupted both discrimination performance and the encoding of stimulus saliency in secondary somatosensory cortical cells (Mo et al., 2024). These findings highlight the importance of transthalamic circuits when considering the neuronal substrates of sensory guided behavior.

In the present study, we tested the transthalamic contribution to perceptual behavior by applying optogenetic inhibition to V1 L5 terminals in the pulvinar in mice during visually guided decision-making. By suppressing V1 L5 inputs to the pulvinar, our approach selectively interrupts transthalamic processing from V1. We found that transthalamic inhibition increased errors and disrupted discriminability across the stimuli presented. These results demonstrate the critical role of L5-initiated transthalamic circuits in visual discrimination behavior.

## RESULTS

### Interrupting L5 Corticothalamic Pathway Originating from Primary Visual Cortex (V1)

We selectively suppressed the axon terminals of V1 L5 projections to the pulvinar using targeted optogenetic inhibition. Expression of the red-shifted inhibitory opsin Jaws (Jaws-tdTomato) (Chuong et al., 2014) was limited to V1 L5 pyramidal cells using Cre-dependent viral injection into V1 of Rbp4-Cre L5 transgenic mice (Gerfen et al., 2013) (Figure 1A). However, as Figure 1A indicates, a large population of L5 neurons of interest in the Rbp-4 mouse does not have Cre (Harris et al., 2014), the implications of which are considered in the *Discussion*. Consistent with previous work, L5 neurons in V1 projected to HO thalamic nuclei — pulvinar and laterodorsal (LD) — and did not project to the first-order visual thalamic nucleus, the lateral geniculate nucleus (Sherman and Guillery, 2013; Usrey and Sherman, 2019; Prasad et al., 2020) (Figure 1B). Red light was delivered to Jaws-expressing V1 L5 terminals in the lateral rostral pulvinar through an implanted optic fiber. This strategy has been shown to inhibit corticothalamic terminal activity in thalamus successfully (Mo et al., 2024).

**Figure 1.**
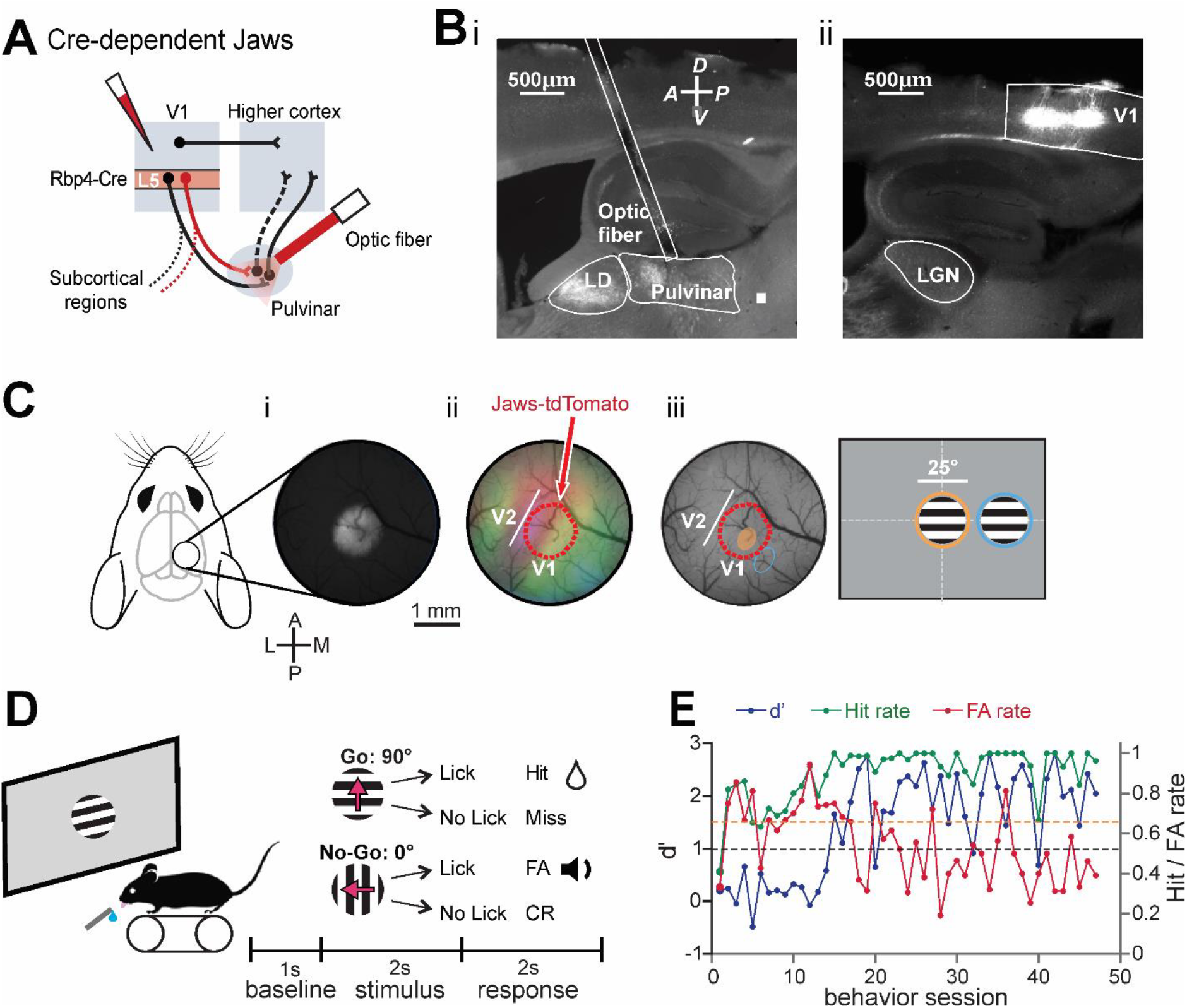
Strategy for inactivation of V1 L5 to pulvinar during visually guided discrimination. **A:** Schematic illustrating strategy for optogenetic inhibition of V1 L5 axon terminals in pulvinar using Cre-dependent Jaws, a red light-driven inhibitory opsin. Note that not all relevant L5 neurons have Cre and thus cannot incorporate Jaws. See text for details. **B**: (i) Sagittal section from an example mouse showing expression of Jaws-tdTomato in V1 L5 terminals located in the pulvinar and LD, as well as the tract from the fiber optic implant. (ii) More lateral section showing Jaws-tdTomato expression in V1 L5 cell bodies and the absence of V1 L5 terminals in the first-order thalamic nucleus, the lateral geniculate nucleus (LGN). A, Anterior; P, posterior; L, lateral; M, medial. **C:** Retinotopic location of Jaws expression in V1 through a cranial window. (i) Jaws-tdTomato fluorescence in cortex. (ii) Retinotopic mapping using IS imaging to functionally determine location of V1 as well as lateral higher visual areas (V2) (Kalatsky and Stryker, 2003). Retinotopic maps overlayed with images of cortical Jaws-tdTomato fluorescence using the vascular patterns as a landmark. The red outline shows the estimated boundary of cortical Jaws expression. (iii) Location of IS imaging responses to task-relevant visual stimuli presented at two positions in the visual field. Monitor schematic shows the relative position of the two stimuli (orange, 0° azimuth, 0° elevation; blue 25° azimuth, 0° elevation). **D:** Schematic of visually guided go/no-go orientation discrimination task. **E:** Plot of behavioral performance for 0° and 90° stimuli across training and testing sessions for an example mouse. The ability to discriminate between orthogonal stimuli improved as a function of training sessions. Orange dashed line indicates d’>1.5 performance criteria. Before beginning testing, mice must perform above this threshold for three consecutive training sessions. Only datasets with performance (d’>1) were included in the analyses.

A cranial window implanted over visual cortex allowed us to identify V1 through intrinsic signal optical (IS) imaging (Kalatsky and Stryker, 2003; Garrett et al., 2014; Juavinett et al., 2017) and to assess the extent of Jaws-tdTomato expression in cortex, thereby determining the retinotopic area of V1 susceptible to L5 terminal suppression (Figure 1C). The visual stimuli in subsequent behavior experiments were retinotopically aligned to the Jaws suppression. The position of the visual stimulus for each mouse was chosen to ensure that the retinotopic location of the visual stimulus response in V1 (determined using IS imaging) fell within the area covered by Jaws expression or, in control experiments, outside that area.

### Visually Guided Go/No-Go Orientation Discrimination Task

Head-fixed mice were trained to visually discriminate between two orthogonally oriented drifting gratings (Figure 1D). The visual stimuli were placed in the hemifield contralateral to the Jaws injections. Using a go/no-go design, the mouse reported responses during a designated response period by licking or withholding licking. Licking in response to the target stimulus (e.g., a 90° or horizontally oriented, upward drifting grating) resulted in a Hit trial, and the mouse was rewarded with a drop of water. A lick following a non-target stimulus (e.g., a 0° or vertically oriented, nasally drifting grating) triggered a false-alarm (FA) trial and was punished with 8 seconds of white noise. For a subset of mice (2 of 5 total), the stimulus reward contingency was reversed (i.e., the go stimulus was vertically oriented). In either case, we refer to the no-go and go stimulus orientations as 0° and 90°, respectively (see *Methods*). Mice were trained to discriminate between 0° and 90° drifting grating stimuli until they reached a high level of discrimination accuracy defined as d’ > 1.5 for three consecutive days (Figure 1E). Mice typically learned the task within 3-5 weeks, at which point they moved on to the testing phase.

### Impaired Visual Discrimination During Inhibition of V1 L5 Projections to Pulvinar

Inhibition of V1 L5 terminals in pulvinar during the stimulus presentation period (Figure 2A) disrupted the ability to discriminate between drifting gratings (0° versus 90°), as indicated by a decrease in d’ (Figure 2B) and an increase in error rate (Figure 2C) for LED-on, relative to LED-off trials. The rise in the error rate was driven by significant increases in both the lapse rate (average lapse rate: LED-off=0.065 ± 0.013, LED-on=0.14 ± 0.026; p=0.012, Wilcoxon signed-rank test, n=25 sessions from 5 mice) and the false alarm rate (average false alarm rate: LED-off=0.34 ± 0.031, LED-on=0.48 ± 0.048; p = 4.7×10^−3^). Notably, optogenetic suppression did not significantly impact the overall rate of lick responses (calculated from hit trials at 90° and FA trials at 0°; p=0.19). This suggests that the observed deficit reflects a decline in perceptual discrimination rather than a more general change in response propensity.

**Figure 2.**
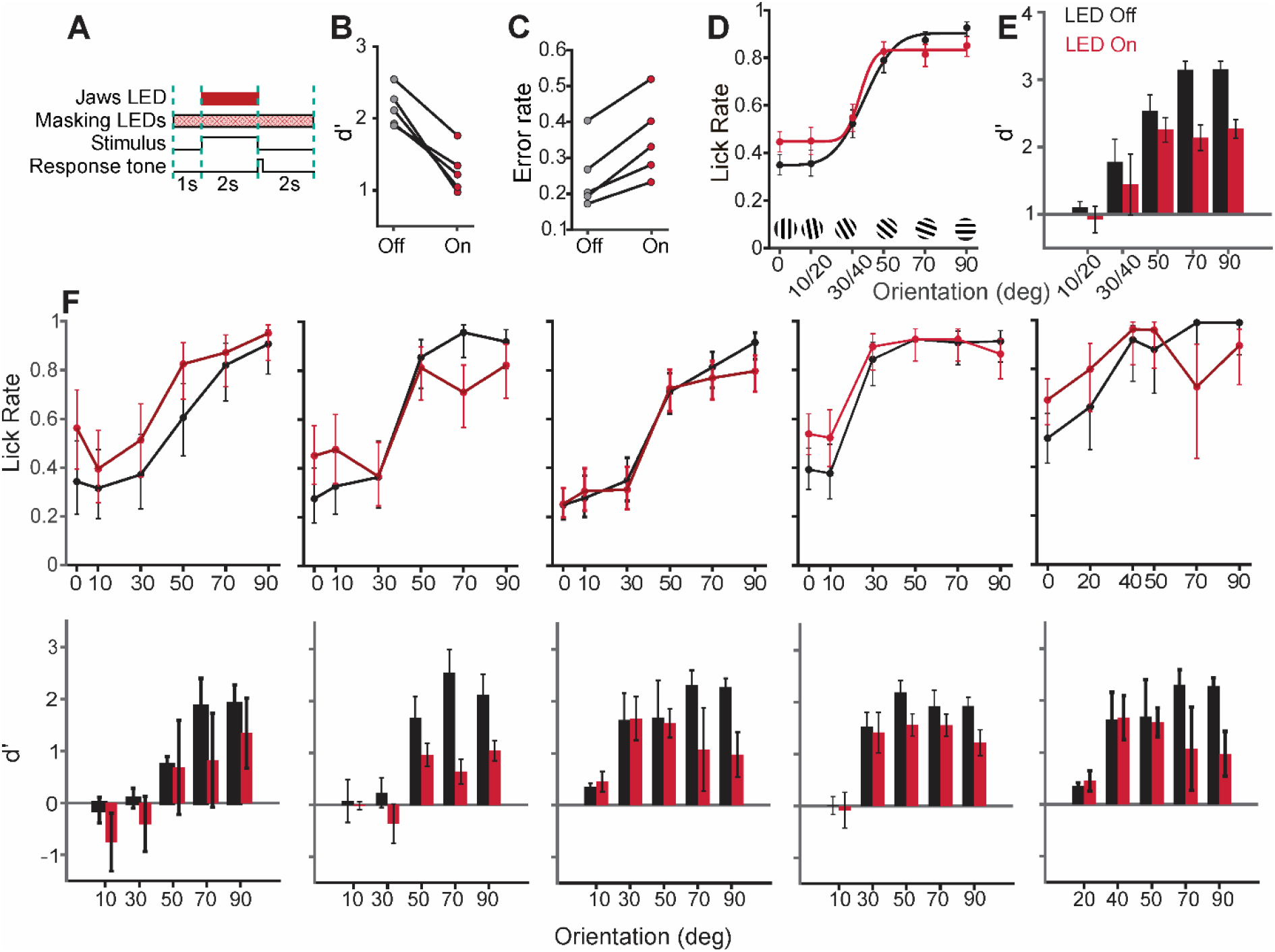
Inactivating V1 L5 projections to pulvinar impairs visual discrimination. **A:** Optogenetic inhibition of V1 L5 terminals in pulvinar during a visually guided discrimination task. For half of the trials, Jaws-expressing V1 L5 pulvinar terminals were inhibited during the sensory presentation epoch by activating a 625nm LED, which delivered red light to the pulvinar via an optic fiber cannula (4.2 ± 0.2 mW at fiber tip). Masking LEDs were used throughout all trials to desensitize the mouse’s retina to red light. **B:** V1 L5 to pulvinar inhibition decreased d’ performance for discriminating 0° and 90° stimuli (average d’: LED-off=2.18 ± 0.14, LED-on=1.32 ± 0.16; p=2.8×10^−4^, Wilcoxon signed-rank test, n=25 sessions from 5 mice). Each pair of data points shows the mean d’ values during LED-off (black) and LED-on (red) conditions for a given mouse (5 mice averaged across 4-7 sessions each). **C**: V1 L5 to pulvinar inhibition induced increase in error rates calculated from FA and Miss trials for 0° and 90° stimuli (average error rate: LED-off=0.24 ± 0.024, LED-on=0.34 ± 0.027; p=9.8×10^−4^, Wilcoxon signed-rank test). **D**: Psychometric curves for LED-on and LED-off trials pooled across all mice (effect of LED x orientation on lick probability: p=3.0×10^−3^, 0°: p=0.018, 10°: p=0.011, two-way RM ANOVA Bonferroni post-hoc test, n=5 mice with 4-7 sessions each). Error bars represent the 95% confidence intervals calculated using a binomial distribution. Note that the abscissa has values of “10/20” and “30/40” because four mice were tested at 10° and 30° and the fifth at 20° and 40° (See **E** and Methods). **E**: Discrimination performance (d’) across stimulus orientations for LED-on (red) and LED-off (black) conditions (effect of LED on d’: p=4.0×10^−3^, effect of LED x orientation: p=0.020, 70°: p=0.022, 90°: p=2.6×10^−3^, two-way RM ANOVA, Bonferroni post-hoc test, n=5 mice with 4-7 sessions each). Error bars indicate mean ± standard error of the mean (SEM) across mice. **F**: Psychometric curves for individual mice from **D**. Trials were pooled across sessions, and error bars show 95% binomial confidence intervals. **G**: Discrimination performance (d’) across stimulus orientations for individual mice from **E** (mean ± SEM across sessions).

In addition to testing discrimination on orthogonal gratings, we assessed psychometric performance by including drifting gratings of orientations varying between 0° and 90°. As expected, in the absence of any experimental manipulation (LED-off), as the angular difference from the “no-lick” (0°) stimulus increased, so did the mouse’s probability of licking (Figure 2D). Jaws suppression appeared to “flatten” the psychometric curve. This change, characterized by a reduction in the slope or “sensitivity” of the curve implies a more homogenous response pattern across stimulus orientations, reflecting a reduced capacity to distinguish between variations in drifting grating orientations. Evidence for this Jaws-induced flattening is seen in each of the individual mice (Figure 2F): For each, the lick rate during Jaws was higher at 0º and, for all but one, lower at 90º.

To further assess the psychometric data, we calculated the discriminability performance using d prime (d’) for all orientations of drifting gratings relative to the no-go (0°) stimulus. Across all stimulus orientations, inhibition significantly reduced d’ values (Figure 2E). The trends of inhibition changing d’ values across stimuli are also evident in data from individual mice (Figure 2G). These findings indicate a significant disruption to visual discrimination performance associated with interrupting processing from V1 L5 to pulvinar.

### No Effect of Red Light When Stimulus Activated Area of V1 Without Jaws Expression

To further control for any unintended effects of the red light used to activate Jaws, we performed the following within-subject control on four of the five mice. During control sessions after expert performance was reached, the experiments were carried out as described above, with the only change being that the stimulus was repositioned on the monitor so that its retinotopic representation in V1 was not within the area affected by Jaws expression (Figure 3A). When the retinotopic location of the stimulus did not correspond with V1 Jaws expression, there was no significant difference in the ability to discriminate orthogonal drifting gratings for LED-on versus LED-off trials (Figure 3B,C). This was also true for psychometric data (Figure 3D, E).

**Figure 3.**
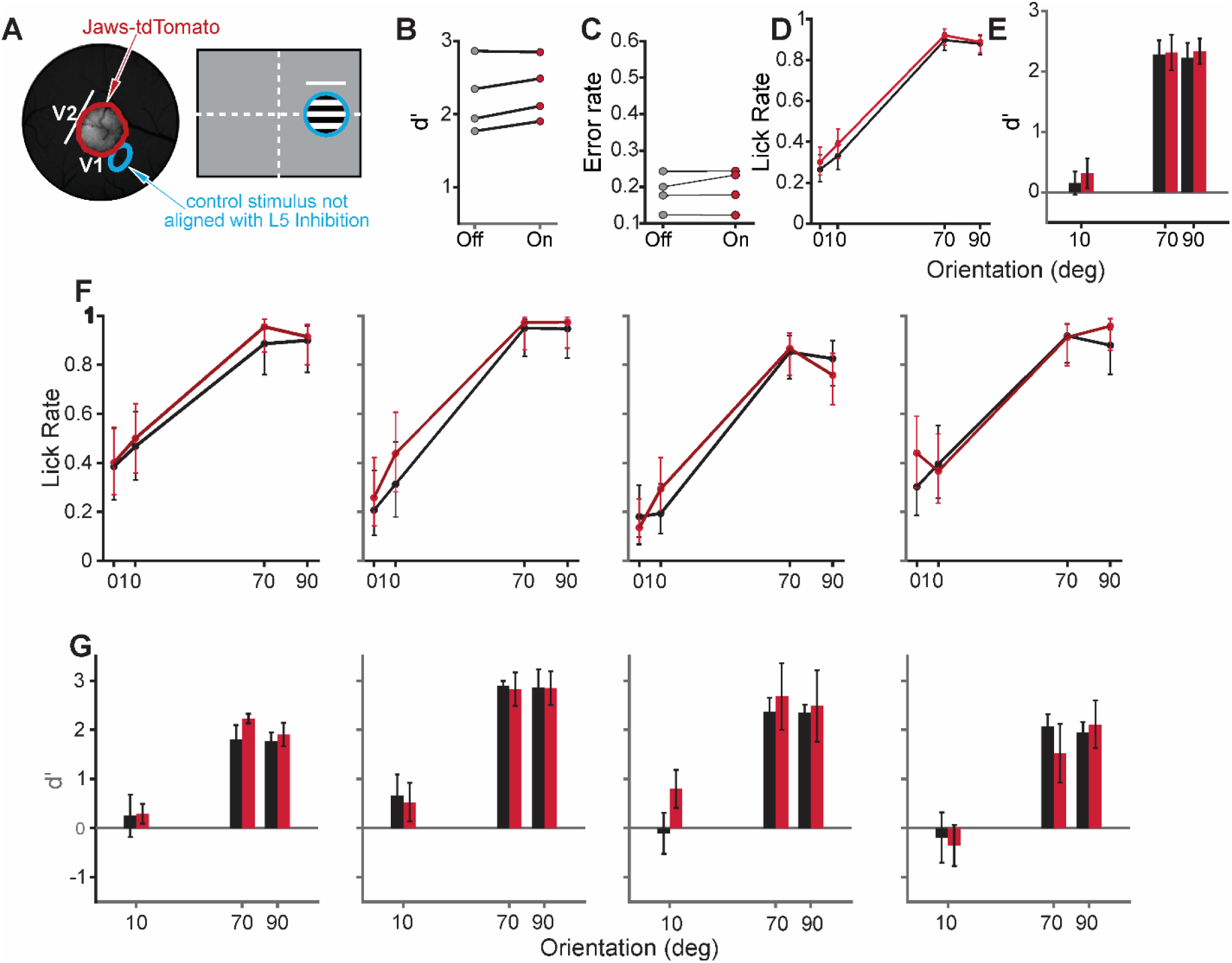
Effects of red light are retinotopically specific to the V1 L5 terminal inhibition. Data from a subset of 4 of the 5 mice shown in Figure 2. **A**: Schematic illustrating within-mouse control experiments. During control experiments, visual stimuli were presented at a position corresponding to a retinotopic location in V1 outside of the area covered by Jaws expression. All other experimental factors remained the same. **B**: Within-mouse control data. There was no effect of the LED (4.2 ± 0.2 mW at fiber tip) on d’ values when the stimulus was located outside of Jaws expression (average d’: LED-off=2.25 ± 0.17 LED-on=2.36 ± 0.23; p=0.59, Wilcoxon signed-rank test, n=14 sessions from 4 mice). **C**: Same as Figure 2C, but for control data (average error rate: LED-off=0.18 ± 0.016, LED-on=0.19 ± 0.026; p=0.79, Wilcoxon signed-rank test). **D**: Similar to Figure 2D. Control data were collected for the four orientations most consistently impacted by inhibition across mice (effect of LED on lick probability: p=0.0.068, two-way RM ANOVA, n=4 mice with 3-4 sessions each). **E**: Same as Figure 2E, but for control data (effect of LED on d’: p=0.73, two-way RM ANOVA). **F**: Individual mouse psychometric data. **G:** Same as in Figure 2G, but for control data. Statistically significant effects of inhibition in the within-Jaws condition remained significant when analyses were restricted to the mice and orientations used in the outside-Jaws condition (0° versus 90° average d’: LED-off=2.17 ± 0.17, LED-on=1.39 ± 0.17; p=1.9×10^−3^, Wilcoxon signed-rank test; 0° vs 90° average error rates: LED-off=0.21 ± 0.021, LED-on=0.31 ± 0.024; p=2.6×10^−3^, Wilcoxon signed-rank test, n=21 sessions from 4 mice).

To ensure an appropriate comparison between the stimulus-within-Jaws and the stimulus-outside-Jaws conditions, we confirmed that statistically significant effects in the within-Jaws experiments persisted when rerunning analyses only included the four mice and four orientations used in the outside-Jaws experiments. The inhibition-induced decrease in d’ and increase in error rates remained significant (Figure 3, control-matched test statistics included at the end of figure legend). Within this sub-sample, data from the within-Jaws experiments was randomly downsampled to match the outside-Jaws sample size to account for differences in the number of sessions between the two conditions (n = 21 within-Jaws and n = 14 outside-Jaws sessions). Across 10,000 iterations of random downsampling, significant effects (p < 0.05) were observed in 98.6% of iterations (mean p = 0.008). Thus, it’s unlikely that the lack of observable effect for outside-Jaws control experiments was due to differences in sampling.

## DISCUSSION

We have demonstrated that suppressing the corticothalamic V1 L5 to pulvinar input during visual discrimination significantly compromises perceptual performance. Specifically, mice exhibited reduced stimulus sensitivity illustrated by shallower psychometric curves. The impact on behavior was observed only when the stimulus representation in V1 was retinotopically aligned with the region affected by Jaws inhibition. Notably, the profound behavioral effect occurred despite our approach primarily affecting neither direct corticocortical projections nor local V1 circuitry; that is, any effects on these are attributed to transthalamic circuits that can effect V1 by a direct input from pulvinar (Miller-Hansen and Sherman, 2022), or by feedback to V1 from other cortical areas innervated by pulvinar. These findings underscore the crucial role of transthalamic corticocortical processing.

### L5 corticothalamic input critical for sensory decision-making

Our findings add to growing evidence that L5 corticothalamic projections to HO thalamic nuclei play a critical role in sensory-based decision-making distinct from that of L5 corticocortical projecting neurons (Takahashi et al., 2020; Qi et al., 2022; Mo et al., 2024). For example, the detection of behaviorally relevant tactile stimuli is determined by the activity of L5 subcortical projection neurons (particularly those to HO thalamus), but not corticocortical projection neurons (Takahashi et al., 2020). Unlike detection, which may not require cortical processing (Sprague, 1966; Schneider, 1969; Sherman, 1974; Hong et al., 2018), discrimination of drifting gratings of differing orientations requires both V1 and higher visual cortical areas, which suggests the hierarchical routing of information between areas (Jin and Glickfeld, 2020). The present study demonstrates the importance of transthalamic processing for successful discrimination of the orientation of drifting gratings. As previously seen during total silencing of V1, the behavioral effects were specific to when the visual stimulus was retinotopically aligned to the optogenetic inhibition, suggesting the function of these pathways is at least in part sensory (Glickfeld et al., 2013; Jin and Glickfeld, 2020). The severe impact of corticothalamic inhibition was seen across the visual gratings presented. This flattening of the stimulus-response curve corroborates with reports of inhibiting cortical L5 to HO thalamic projections in a somatosensory discrimination task (Mo et al., 2024). These behavioral data suggest that this corticothalamic projection, and thus transthalamic circuitry, is necessary for correct perceptual discrimination.

### Potential downstream pulvinar targets

By inhibiting V1 L5 driving input to pulvinar, we interrupted the first synapse of di-synaptic transthalamic circuits and, therefore, disrupted signal transmission to downstream cortical targets as well as V1 itself. Pulvinar outputs widely target visual cortical areas (Bennett et al., 2019; Juavinett et al., 2020; Mease and Gonzalez, 2021). Based on these thalamocortical outputs, there are two established connectivity patterns through which transthalamic circuits originating from V1 L5 impact cortical processing.

#### Recurrent loops back to V1

Some V1 L5 inputs drive pulvinar cells that send modulatory projections back to V1 (Bennett et al., 2019; Miller-Hansen and Sherman, 2022; Cassidy et al., 2024). Pulvinar input to L1 may provide V1 sensory signals with contextual modulation related to the animal’s motor outputs, such as increasing the salience of visual motion that is not due to the movement of the animal itself (Roth et al., 2016). However, recent work suggests that feedback transthalamic circuits that convey strong driving input throughout, which would violate the “no-strong-loops” hypothesis (Crick and Koch, 1998), are rare or nonexistent (Bennett et al., 2019; Miller-Hansen and Sherman, 2022; Cassidy et al., 2024).

#### Feedforward pathways to higher cortical areas

V1 L5 projections also synapse onto extrastriate-projecting pulvinar neurons, initiating feedforward transthalamic circuits (Blot et al., 2021; Miller-Hansen and Sherman, 2022). In contrast to pulvinar projections to V1, pulvinar projections to extrastriate cortical areas provide substantial driving input that heavily influences cortical responses (Zhou et al., 2016; Zhou et al., 2018; Beltramo and Scanziani, 2019; Miller-Hansen and Sherman, 2022). This is consistent with the idea that feedforward transthalamic pathways provide powerful indirect routes for information transfer between primary and higher cortical areas (Theyel et al., 2010; Mo et al., 2024). The pulvinar also sends projections to frontal and parietal cortical regions, including the anterior cingulate cortex (Kaas and Lyon, 2007; Bennett et al., 2019), which may be additional candidates for feedforward transthalamic input from visual cortex (Leow et al., 2022).

#### Noncortical targets

Although nearly all pulvinar outputs target cortex, some of its projections also send branches to the striatum and amygdala (Day-Brown et al., 2010; Wei et al., 2015; Bennett et al., 2019). Projections to these subcortical structures arise from superior colliculus-recipient regions of pulvinar with minimal overlap with cortical L5 projections, making it unlikely that the main transthalamic targets relative to our findings are subcortical (Day-Brown et al., 2010; Zhou et al., 2017). Nonetheless, our data do not allow us to identify which downstream areas, targeted by V1 L5-recipient pulvinar projections, are impacted in our experiments. Future studies are needed to address the specific pulvinar targets involved.

### Transthalamic circuits may signal the behavioral relevance of sensory stimuli

Recent evidence supports the idea that feedforward transthalamic circuits contribute to sensory decision-making by establishing task-dependent cellular selectivity in higher cortical areas (Yang et al., 2022). In mice performing a somatosensory-based discrimination task, thalamic input, not the direct corticocortical input, drives stimulus and choice selectivity in frontal cortex (Yang et al., 2022). Consistent with this, inhibiting somatosensory L5 corticothalamic input impaired discrimination performance and interrupted reward stimulus encoding in higher order cortex (Mo et al., 2024). The inhibition also had a more disruptive effect on representations in secondary somatosensory cortex relative to primary (Mo et al., 2024), consistent with the importance of the feedforward pathway to higher order cortex. This suggests that the transthalamic pathway carries task-relevant information up the cortical hierarchy. In the visual system, transthalamic pathways through pulvinar are thought to distinguish between self-generated signals and external visual stimuli, thus playing a role in integrating visual information with the broader behavioral context (Blot et al., 2021). Additionally, findings from non-human primates suggest that pulvinar-mediated transthalamic circuits coordinate interactions between cortical areas based on task demands, allowing for more efficient transmission of behaviorally relevant sensory information (Saalmann et al., 2012; Halassa and Kastner, 2017; Jaramillo et al., 2019).

### Some qualifications for our data interpretation

*V1 L5 transthalamic circuit through LD*. We have concentrated above on transthalamic circuits involving pulvinar. However, in addition to projecting to the pulvinar, Figure 1 shows that L5 neurons from V1 also send projections to the HO thalamic nucleus, LD (Prasad et al., 2020). Like pulvinar, mouse LD sends projections to all visual cortical areas (Juavinett et al., 2020), and suppression of V1 L5 projections to LD would similarly disrupt transthalamic pathways originating from V1. However, the likelihood of suppressing LD terminals is low, given the distance and expected irradiance loss. Specifically, Jaws-expressing terminals in LD are located more than 500 µm from the tip of the optic fiber, a distance at which over 50% of irradiance is lost (see http://www.stanford.edu/group/dlab/cgi-bin/graph/chart.php). This exponential loss in light power is also a sound argument against any direct effects of inhibiting V1 L5 cells, which sit at a distance from the optic fiber where irradiance is negligible. Additionally, for LD, the posterior-facing angle of the optic fiber implant further reduces the probability that light delivered to the pulvinar would affect the more anteriorly located LD. Nonetheless, even if circuitry via LD were responsible for some of our data, it would not affect our main conclusion that transthalamic circuitry starting from L5 of V1 is necessary for visual discrimination.

#### Uncertainty about eye position

One potential limitation of our study is the absence of eye-tracking data to precisely determine the mouse’s eye position. However, this is largely mitigated by the observation that, in mice, eye movements are tightly linked to head movement (Meyer et al., 2020; Michaiel et al., 2020). To maintain consistency of retinotopic mapping, the mouse’s head position relative to the monitor was kept constant across IS imaging and behavior sessions. While eye movement within this head-fixed preparation cannot be completely ruled out (Meyer et al., 2020), our results indicate that behavioral changes were dependent on the stimulus being retinotopically aligned with the inhibition, providing compelling empirical evidence that eye movements did not distort the retinotopic targeting of visual stimuli.

#### Underestimating effect

Our findings almost certainly strongly underestimate the impact of inhibiting V1 L5 projections to pulvinar. This is because our approach could not have affected all, or even most L5 terminals in pulvinar, for two main reasons: 1) We inserted Jaws into these L5 terminals using a Cre strategy, but many such L5 cells do not express Cre in our transgenic mice (Harris et al., 2014). 2) We do not expect all the Cre-positive L5 terminals transfected with Jaws to have been entirely inhibited by our light probe activation.

### Conclusions

We demonstrated that V1 L5 projections to pulvinar are critical for visual discrimination. Our findings in the visual system align with recent evidence from the somatosensory system showing that L5 corticothalamic input to HO thalamic nuclei is critical for stimulus detection and discrimination (Takahashi et al., 2020; Qi et al., 2022; Mo et al., 2024). The potential for similar findings across sensory systems highlights the importance of studying L5 corticothalamic inputs and their role in sensory processing and decision-making. The findings of such work have broad implications for our understanding of how information propagates through the cortex.

## MATERIALS AND METHODS

### Animals

All procedures were approved by the Institutional Animal Care and Use Committee at the University of Chicago. To breed transgenic mice with Cre recombinase expression in cortical L5 (Gerfen et al., 2013), female C57BL6J mice were crossed with male Tg(Rbp4-Cre) KL100Gsat/Mmcd mice (GENSAT RP24-285K21). Tail biopsies were taken at 14-21 days old and genotyped by real-time polymerase chain reaction (Transnetyx, Cordova, TN). Data were obtained from 5 adult Rbp4-Cre positive mice of both sexes (3 male, 2 female). Mice were housed individually on a 12-hour reverse light/dark cycle (dark from 7am-7pm), with food and water freely available except during water restriction as described.

### Surgical Procedures

Surgery procedures followed those previously described (Mo et al., 2024) with modifications for the visual system. Briefly, before surgery, mice were anesthetized by intraperitoneal injection of ketamine (100 mg/kg)/xylazine (3 mg/kg) and then maintained under anesthesia with isoflurane (1.0-1.5% in oxygen). While under anesthetic, a heating pad was used to maintain body temperature, and lubricant was used to prevent the eyes from drying. To achieve viral expression of Jaws-TdTomato in L5 of V1, 9-12-week-old mice were injected with AAV8-CAG-FLEX-Jaws-KGC-TdTomato-ER2 (UNC Vector Core) in the left hemisphere V1 (coordinates from bregma, DV: −0.5mm, ML: 2.5mm, AP: -4.0mm) using a 0.5 uL Hamilton syringe (300 nl at 10-15 nl/min). Following a two-week recovery period, a custom head plate was cemented to the skull, a 4mm cranial window was implanted over the left visual cortex (center of window relative to lambda, ML: 4.1mm; AP: 0.5mm), and a fiber optic cannula (see *Optogenetic Inhibition*) was implanted in the left lateral rostral pulvinar. The optic fiber was implanted at a 25° angle anterior from vertical to accommodate the cranial window. When the stereotaxic instrument was at a 25° anterior angle, the coordinates for implantation were (DV: -2.8mm, ML: 1.6mm, AP: -0.8mm).

### Intrinsic Signal Optical Imaging

Intrinsic signal (IS) optical imaging was performed using a previously described setup (Mo et al., 2024). Anesthetized mice (induction: 3% isoflurane in oxygen, maintenance: 1% isoflurane) were head-fixed under a CCD camera (Teledyne QImaging, Retiga-SRV), and the intrinsic hemodynamic signal (measured by changes in 625nm light reflectance) was imaged through the cranial window. The cortical surface vasculature was visualized under green (525nm) light illumination.

Visual stimuli were presented to the eye contralateral to the imaged cortex (Dell P2412Hb). Images were acquired using custom MATLAB code, and stimulus presentation was performed using custom MATLAB code in combination with Psychophysics Toolbox (Brainard, 1997). IS imaging was first performed while presenting a continuous moving bar stimulus to map cortical retinotopy (Kalatsky and Stryker, 2003) and functionally locate V1. Next, we looked at V1 responses to the behavior-relevant stimuli at various locations in the visual field. Stimuli consisted of upward drifting sinusoidal gratings (2 Hz; 0.04 cycles/degree) presented through a 25° diameter circular aperture on a mean luminance-matched grey background. Each imaging trial began with 6 seconds of a mean luminance-matched grey screen, followed by 10 seconds of stimulus presentation. Stimulus response was calculated as the percent change in reflected light between the average of the last 4 seconds of the baseline period and the average of the first 8 seconds of the stimulus period.

### Visual Discrimination Task

During behavioral sessions, mice were head-fixed and could run freely on a custom-built treadmill. Based on a go/no-go design, the mouse reported responses during the designated response period by licking or withholding licking. The waterspout (14G blunt needle) was positioned 3-6 mm from the mouth, and licks were detected using a capacitance sensor (Teensy 3.2, PJRC). On rewarded trials, water (8µl) was dispensed through the spout via a pump (NE-1000 syringe pump, New Era). Behavioral training and testing were implemented with custom MATLAB code using the Psychophysics Toolbox (Brainard, 1997) for visual stimulus generation. Stimuli were presented to the mouse on a monitor (ASUS VG248) positioned 20cm from the right eye. Stimuli consisted of drifting sinusoidal gratings (2 Hz, 0.04 cycles/degree) presented through a 25° diameter circular aperture on a mean luminance-matched grey background. A strip of 13 625nm LEDs was mounted below the monitor. The behavioral setup was housed inside a lightproof enclosure lined with soundproof material (0.8 NRC, Sound Seal). An infrared webcam was used to monitor inside the enclosure.

Mice were trained to discriminate between two orthogonally oriented drifting gratings. Licking in response to the target stimulus (90°, e.g., horizontally oriented upward drifting grating) was classified as a hit trial, and the mouse was rewarded with a drop of water. A lick following a non-target stimulus (0° e.g., vertically oriented rightward drifting grating) was classified as a false-alarm (FA) trial and was punished with 8 seconds of white noise. Failing to lick in response to the target stimulus was classified as a miss, whereas successfully withholding licking in response to the non-target stimulus was regarded as a correct rejection (CR). The stimulus reward contingency was reversed for two of the five mice (i.e. the target was vertically oriented). For consistency across animals, we refer to the non-target stimulus orientation as 0° and the orthogonal target stimulus as 90° (indicating the angular distance from the non-target).

Following at least one week of post-surgery recovery, mice were water-restricted to 80-95% of their body weight and habituated to head fixation. During habituation sessions, water was freely delivered through the spout to encourage licking. Following this, water delivery was associated with a preceding tone (12kHz, 60dB) until the mice learned to self-initiate the water release by licking the spout within 2 seconds of hearing the tone. Once the mouse could reliably lick following the response tone (typically 3-5 days from the start of habituation), they moved on to the discrimination training phase.

In the first phase of discrimination training, mice were familiarized with the trial structure and learned to associate the target stimulus with the reward. The stimulus monitor remained black during intertrial intervals (3-9 seconds). A blank grey screen (with mean luminance matched to the gratings) indicated the start of a trial. After a 1 second baseline period, the target stimulus was presented for 2 seconds. At the end of the stimulus period, the monitor returned to grey, and a response tone (12kHz, 60dB) indicated the start of the response period (2 seconds). The water reward was automatically dispensed immediately following the response tone for the first few trials. Thereafter, mice received the reward only if they licked during the response window (hit). Licks that occurred outside of the response period were ignored. Once the mice reliably licked in response to the target stimulus, the non-target (0°) stimulus was introduced. In cases of excessive licking during non-target stimulus trials, the FA white noise punishment was accompanied by a short air puff to the snout (Cleaning duster, Office Depot).

During initial training, the stimulus size was increased to fill the full height of the monitor (60° diameter) and was centered in the middle of the screen. Once the mouse could reliably discriminate between the 0° and 90° oriented drifting gratings, we gradually shrunk and shifted the stimulus until it was 25° degrees in diameter and centered on the mouse-specific coordinates (see *Intrinsic signal optical imaging*). Mice were trained on 0° versus 90° discrimination until they reached a performance criterion of d’ > 1.5 for three consecutive days, at which point they moved on to the testing phase.

Each experimental session began with a “warm-up” period to mitigate any initial bias toward go responses (Berditchevskaia et al., 2016). During this phase, mice performed 0° versus 90° discrimination trials, and testing was initiated only after they correctly rejected (CR) the no-go stimulus three consecutive times (with go-stimulus trials randomly interleaved). To collect psychometric data during behavioral testing, stimuli included gratings of various orientations ranging between 0° and 90° (0°, 10°, 30°, 50°, 70°, 90° for four mice, and 0°, 20°, 40°, 50°, 70°, 90° for the fifth mouse). Stimuli were presented pseudo-randomly but with no more than three of one type in a row. To further discourage a predisposition for lick responses, the non-target stimulus (0°) was presented to 3 of the 5 mice in a 2:1 ratio relative to other stimulus orientations. Both 0° and 10° were treated as non-target stimuli for the other two mice, and all orientations were presented in equal proportions. If mice showed perseveration (licking at every trial) or disengagement (no lick responses), they reverted to 0° versus 90° training (Mo et al., 2024). All 0° versus 90° bias correction trials (i.e. warm-up and retraining) were excluded prior to analyses, as detailed below. Behavior sessions were run 5-7 days/week, each lasting 1-2 hours (∼200 trials). Data from all behavioral sessions were preprocessed post hoc using an automated MATLAB-based pipeline to crop task-relevant trials, excluding non-representative trials such as the warm-up and retraining phases. The predefined criteria were as follows. (1) The algorithm identified session onset as the first trial meeting test conditions (i.e LED-on or stimulus at an orientation other than 0° or 90°) following the end of the warm-up period, marked by three consecutive CRs without interleaved misses. All trials before this point were excluded. (2) Retraining periods, defined as > 4 consecutive LED-off trials at 0° or 90°, were flagged and removed along with a buffer of 10 preceding trials. (3) Sessions were terminated at the first LED-off trial at which one of the following criteria was met, indicating disengagement or satiation: (3a) three consecutive misses at 70° or 90° (easiest to discriminate from 0°); (3b) a hit rate < 0.40 (calculated over a 40 trial slide window); (3c) an overall lick rate (across all stimulus orientations) > 0.90. This automated trial cropping approach ensured a consistent, unbiased data selection for subsequent analyses. Analyses included only the sessions where mice maintained a 0° versus 90° discrimination performance of d’ > 1 for trials without optogenetic manipulation.

### Optogenetic Inhibition

Optogenetic inhibition was performed using the inhibitory opsin Jaws, a red-light sensitive chloride pump (Chuong et al., 2014). Light from a 625nm LED (Thorlabs) was delivered through a patch cable (0.5NA, Thorlabs) to the implanted optic fiber (200µm diameter, 0.5NA, Thorlabs), which then illuminated Jaws-expressing V1 L5 terminals in the pulvinar. One of the five mice received red light through a 0.39NA optic fiber (200µm diameter, 39NA, Thorlabs) and patch cable (0.39NA, Thorlabs). In all cases, the estimated power output at the fiber tip was 4.2 ± 0.2 mW (∼134 mW/mm^2^). The LED was turned on during the stimulus period (2 seconds) for 50% of trials. To minimize the possibility that mice could perceive the red light used to activate Jaws, the retina was habituated to 625nm light via a strip of masking LEDs positioned under the stimulus monitor (Danskin et al., 2015; Odoemene et al., 2018), which remained on for the session.

During testing, the position of the task stimulus for each mouse was chosen so that the stimulus representation in V1 (see *Intrinsic Signal Optical Imaging*) retinotopically aligned with the L5 to pulvinar projection inhibition. When collecting within mouse control data, the experiments were carried out as described above except that the stimulus was repositioned on the monitor so that its retinotopic representation in V1 was not within the area affected by Jaws expression.

### Behavioral Data Analyses

Discrimination performance was quantified using d’, calculated as the difference between the normalized Hit and FA rates:

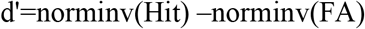

where norminv() is the cumulative normal function. Hit and FA rates were cut off at 0.99 and 0.01, giving a maximum possible d’ of 4.65. Higher d’ values indicate that the animal is more likely to lick in response to the “lick” stimulus and less likely to respond to the “no-lick” stimulus, reflecting better discrimination.

Error rate was calculated from 0° and 90° trials as follows:

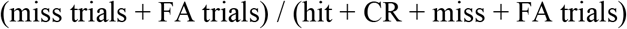

Psychometric curves were fitted to response rates with a 4-parameter sigmoidal cumulative Gaussian function (Wichmann and Hill, 2001):

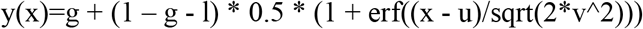

where y(x) is the lick probability, x is the orientation, and erf is the error function. The parameters to be fitted are g (guess rate), l (lapse rate), u (subject bias), and v (discrimination sensitivity).

### Fluorescence Microscopy

Mice were perfused with cold 0.1 M phosphate-buffered saline (PBS) followed by cold 4% paraformaldehyde (PFA). Each brain was extracted and post-fixed in 4% PFA overnight and then stored in 30% sucrose in PBS for 2 days. Using a sliding microtome, the brain was sectioned into 50µm thick sagittal slices and mounted on Superfrost slides. Fluorescence signals were visualized under a fluorescence microscope (Leica Microsystems) using the appropriate filter cubes. Images were captured using a Retiga-2000 CCD monochrome camera and QCapturePro imaging software (Teledyne QImaging, Surrey, BC). Image post-processing was performed with ImageJ software.

### Statistics

Statistical comparisons were performed in MATLAB. Sample sizes, statistical tests, and associated p-values are included in the figure legends. Average values are reported as mean ± sem. Paired samples were analyzed using the Wilcoxon signed-rank test. A two-way repeated measures ANOVA was used to investigate the effects of LED condition and stimulus orientation. The Geisser-Greenhouse correction was applied to adjust for deviations from sphericity. Our significance threshold of p<0.05 was adjusted for multiple comparisons using Bonferroni correction.

## Data Availability

Data is available from the corresponding author upon reasonable request.

## ACKNOWLEDGMENTS

This work was supported by the National Institutes of Health (Grants NS094184 and EY036242 to S.M.S. and F31EY031965 to C.McK.) and the National Health and Medical Research Council, Australia (Grant 2003646 to C.M.). We thank Adam Kunz for his help with behavioral training.

